# cancerSimCraft: A Multi-resolution Cancer Genome Simulator with Comprehensive Ground Truth Tracking

**DOI:** 10.1101/2024.12.11.627708

**Authors:** Haijing Jin, Nicholas Navin, Ken Chen

## Abstract

Cancer evolution follows complex trajectories involving diverse genomic alterations and clonal dynamics, making it challenging to validate computational methods for single-cell DNA sequencing analysis. Here we present cancerSimCraft, a comprehensive framework for simulating cancer genome data at both clonal and single-cell resolution. cancerSimCraft combines deterministic rules with stochastic processes to model various genomic events including CNVs, SNVs, and WGDs. The framework enables integration of real cancer genome patterns with user-defined parameters, supporting customizable simulation designs that reflect both empirical data and theoretical models. Through systematic benchmarking, we demonstrate cancerSimCraft’s utility in evaluating computational methods under various conditions, particularly focusing on the impact of dataset size and parameter sensitivity. We further demonstrated its application in exploring clonal evolution and mutation patterns through controlled *in silico* experiments. These comprehensive simulation capabilities make cancerSimCraft a valuable resource for both computational method development and theoretical studies in cancer genomics.

## Introduction

Cancer is a complex disease characterized by the uncontrolled growth and proliferation of somatic cells (Hanahan & Weinberg, 2011). Cancer develops through an evolutionary process where somatic cells acquire advantageous mutations under selective pressures, leading to the emergence of distinct subclones (Shendure & Akey, 2015; Tarabichi et al., 2021). The interplay of random mutations and natural selection through evolutionary processes drives extensive tumor heterogeneity, complicating effective diagnosis and treatment (Fisher et al., 2013; Zhu et al., 2021).

Recent technological advancements in scDNA-seq technology have revolutionized cancer research by enabling the detection of genetic alterations with unprecedented resolution (Jia et al., 2022). While bulk DNA-seq data provides averaged genetic signals across heterogenous cell populations in tissue samples, scDNA-seq data captures the genetic information from individual cells, preserving the inherent cellular diversity within a tumor. This high-resolution, single-cell approach enables the detection of rare cancer subclones, elucidates clonal relationships, and reveals complex evolutionary trajectories that are often obscured in bulk sequencing data (Minussi et al., 2021; Pang et al., 2024). Through the characterization of fine-scale genetic variations, scDNA-seq enhances our fundamental understanding of tumor heterogeneity and clonal evolution while providing critical insights into clinically relevant phenomena such as therapy resistance mechanisms and treatment response prediction (Demaree et al., 2021; Kim et al., 2018).

In recent years, numerous computational methods have been developed to analyze single-cell DNA sequencing (scDNA-seq) data for cancer clonal reconstruction. The methodological spectrum encompasses a diverse set of analytical tools: Copykit (Minussi et al., 2022) for total copy number (CN) determination, CHISEL (Zaccaria & Raphael, 2021) for haplotype copy number analysis, MEDICC2 (Kaufmann et al., 2022) for phylogenetic inference, and Monopogen (Dou et al., 2024) for variant characterization. Despite these advancements, evaluating accuracy and reliability of these methods remains a significant challenge. Current validations rely predominantly on benchmarks presented within original publications of computational tools (Dou et al., 2024; Kaufmann et al., 2022; Minussi et al., 2022; Zaccaria & Raphael, 2021), where evaluation criteria and dataset selection are inherently inconsistent across studies. This absence of standardized and systematic benchmarking frameworks precludes objective performance comparisons across different experimental and computational settings.

To address this limitation, researchers have launched a crow-sourced benchmarking challenge – the ICGC-TCGA DREAM Somatic Mutation Calling Tumor Heterogeneity Challenge (SMC – Het) (Salcedo et al., 2020). This initiative facilitates the objective comparison of computational tools by employing standardized simulated tumor datasets and scoring metrics. However, the SMC-Het benchmark predominately relies on bulk sample simulations, which lack the resolution to fully capture the complexity and heterogeneity of single-cell data. To overcome this limitation, we need simulators that can achieve single-cell and single-nucleotide resolution simulations with comprehensive ground truth tracking.

Beyond developing simulators that can mimic genetic alterations at single-cell resolution, it is essential to establish a streamlined workflow that integrates simulation design, data generation, and benchmarking procedures. Such an approach would enable research groups to create customized simulation frameworks for their specific research needs. Given that cancer arises from different tissues and patient populations (Melo et al., 2013; Seferbekova et al., 2023), and follows distinct evolutionary trajectories (Vendramin et al., 2021), adaptable simulators are crucial for capturing these diverse biological characteristics.

To address these challenges – particularly the need for high-resolution, customizable simulations – we developed cancerSimCraft, a cancer genome simulator that combines deterministic rules and stochastic processes to derive simulations at single-cell and single-nucleotide resolution at the whole-genome scale. cancerSimCraft allows researchers to integrate information from multiple sources, including biological knowledge, real cancer genome data, and user-defined hypotheses, supporting realistic simulations that reflect the intricacies of cancer genomes. Through its modular architecture, cancerSimCraft separates simulation design from the data generation process, enabling efficient parameter exploration without redundant computational processing. Leveraging parallel computing to support multi-core processing, cancerSimCraft achieves substantially higher throughput than existing simulation methods (Mallory & Nakhleh, 2022) in single-cell resolution simulation, making it well-suited for high-throughput applications in cancer genomics research. A key feature of cancerSimCraft is its comprehensive ground-truth tracking system, which records every simulated genomic alteration at both clonal and single-cell levels. This comprehensive tracking system enables both customizable benchmarking of computational workflows and systematic *in silico* experimentation, allowing researchers to validate methods and explore hypotheses through simulation.

A series of systematic analyses have been conducted to comprehensively evaluate cancerSimCraft’s utility and reliability. We first validated cancerSimCraft’s simulation reliability by designing rule-based simulations that recapitulate key features of real cancer genomes, followed by comprehensive validation at both genome and read levels. We then demonstrated its applications in benchmarking computational methods for cancer genome analysis, particularly focusing on subclone assignment and copy number inference. Finally, we illustrated how cancerSimCraft can be used to explore theoretical questions through systematic *in silico* experiments. Taken together, these analyses demonstrate cancerSimCraft as a robust and versatile platform for cancer genome research, supporting diverse applications such as realistic simulation, experimental design, benchmarking of computational tools, and *in silico* exploration of theories.

## Results

### Overview of cancerSimCraft’s architecture and applications

We present cancerSimCraft, a comprehensive genome simulator designed to computationally synthesize cancer genomes with flexibility, precision, and scalability. cancerSimCraft is designed to integrate multiple sources of information, including curated biological knowledge, cancer genome data, and user-defined hypotheses (Fig. 1a). This integration enables researchers to simulate cancer genomes that reflect not only signal patterns from real data but also biological insights, providing a foundation for experimental design, benchmarking, and *in silico* experimentation (Fig. 1a).

**Figure 1.**
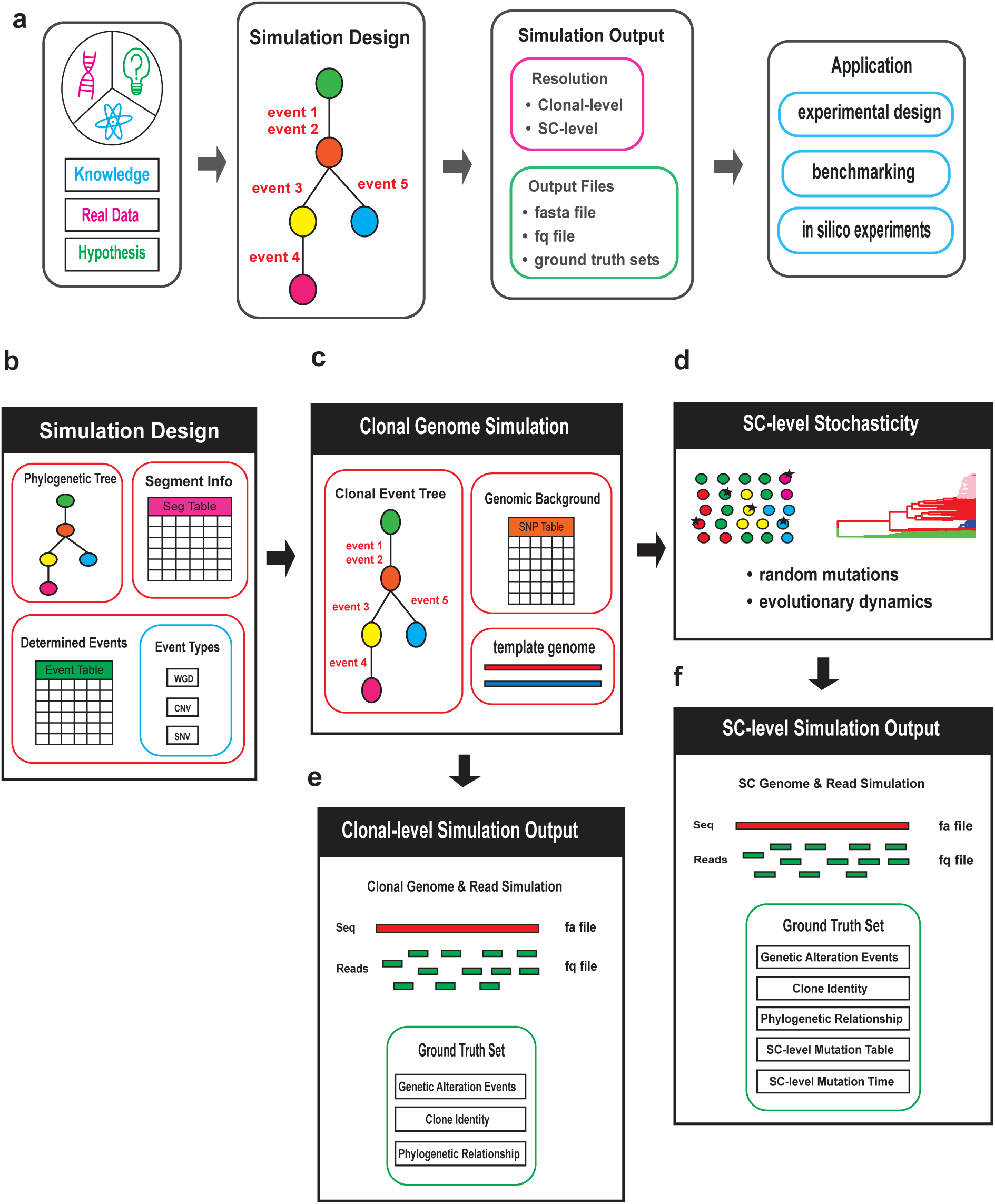
Framework and implementation of cancerSimCraft. **a,** Overview of cancerSimCraft’s framework: The simulator integrates information from multiple sources, including knowledge, real cancer genome data, and user-defined hypotheses. Its modular architecture separates simulation configuration from execution, enabling efficient customization of simulation settings. The simulator outputs standard sequencing data formats (FASTA, FASTQ) and comprehensive ground truth records. These features make cancerSimCraft a powerful tool for various applications, such as experimental design evaluation, computation tool benchmarking, and systematic *in silico* exploratory studies. **b,** Core components of the simulation design: The simulator requires three primary inputs: (1) a pre-defined phylogenetic tree describing the evolutionary relationships between cell populations, (2) an event table specifying genomic alterations, and (3) a segment table defining genomic regions of interest. The framework supports multiple types of genomic alterations, including single nucleotide variants (SNVs), copy number variations (CNVs), and whole genome duplication (WGD) events. **c,** Genome simulation process: The simulator first creates a clonal event tree by integrating phylogenetic relationships with subclone-specific genomic alterations. Users can optionally incorporate patient-specific variation from a SNP table. Using a template genome (typically the human reference genome) as the base, cancerSimCraft generates unique genomes for each subclone in accordance with the preset evolutionary relationships and genomic alterations. **d,** Single-cell stochastic modeling: To simulate evolutionary dynamics at the single-cell level, cancerSimCraft employs a continuous-time Markov process to model discrete cellular and genomic events, including cell birth, death, transformation, and mutations. This stochastic layer introduces realistic cell-to-cell variation, capturing both cellular dynamics and genomic alterations to reflect the complexity of cancer progression. **e,** Clonal-level simulation outputs: The simulator produces standard genome file formats (fasta and fastq) alongside comprehensive ground truth data. The ground truth includes details on genomic alterations, clone identities, and the phylogenetic relationships of all simulated read files (fastq). Each subclone’s simulated read files are generated based on a shared subclone-specific genome. **f,** Single-cell level simulation outputs: Similar to clonal-level simulation, cancerSimCraft generates both fasta and fastq files and ground truth data for single-cell simulation. In addition to clonal-level information, the ground truth includes single-cell mutation tables and mutation timing data. Each simulated fastq file represents a unique single-cell genome, incorporating both clonal-level and single-cell-specific alterations.

A key feature of cancerSimCraft is its modular architecture, which separates simulation configuration from execution (Fig. 1a). This allows researchers to adjust simulation settings independently without rerunning the entire simulation, enabling efficient customization and reuse of configurations. With decoupled design and execution processes, cancerSimCraft facilitates rapid iteration and testing of different simulation scenarios, enhancing both the scalability and flexibility of the tool.

In terms of output, cancerSimCraft generates data in standard sequencing formats, including fasta and fastq, which are widely compatible with most bioinformatics tools (Fig. 1a). Alongside these files, the simulator produces detailed ground truth that documents every simulated genomic alteration and cellular event. This comprehensive tracking of ground truth enables direct comparison between simulated data and computational predictions or experimental results, making cancerSimCraft a well-suited tool for method validation and benchmarking. Moreover, cancerSimCraft supports systematic exploratory studies in cancer research, enabling researchers to test hypotheses and explore complex genomic and cellular dynamics in a virtual setting.

### Multi-Level cancer genome simulation and ground truth tracking in cancerSimCraft

To achieve precise cancer genome simulation, cancerSimCraft employs a systematic workflow that incorporates both deterministic and stochastic elements. Its core architecture relies on three fundamental inputs (Fig. 1b): a phylogenetic tree defining evolutionary relationships between cell populations, an event table specifying genomic alterations, and a segment table that divides the genome into predefined regions (bins). The bin sizes can be customized based on the analysis requirements. The simulator supports various types of genomic alterations, including single nucleotide variants (SNVs), copy number variations (CNVs), and whole genome duplication (WGD) events. After initial configuration, the genome simulation module in cancerSimCraft combines these inputs into a clonal event tree, which integrates phylogenetic relationships with subclone-specific genomic alterations (Fig. 1c and Supplementary Fig. 1b). Researchers can also incorporate patient-specific variation through an optional SNP table. Using a template genome (typically the human reference genome) as a base, cancerSimCraft generates unique genomes for each subclone in accordance with the predefined simulation configuration. A key innovation of cancerSimCraft lies in its ability to integrate single-cell evolutionary modeling with a detailed ground truth tracking system. The simulator’s ability to model single-cell dynamics through a continuous Markov process (Fig. 1d) allows it to simulate various cellular events including birth, death, transformation, and mutation accumulation, reflecting the inherent heterogeneity derived from cancer evolution at single-cell resolution (Methods).

The simulation outputs of cancerSimCraft are produced at both the clonal (Fig. 1e) and single-cell (Fig. 1f) levels. At the clonal level, cancerSimCraft generates standard sequence files (fasta and fastq) along with comprehensive ground truth data covering genomic alterations, clone identities, and phylogenetic relationships across all simulated samples (Fig. 1e). At the single-cell level, each simulated fastq file represents a unique single-cell genome that captures both clonal-level alterations and single-cell-specific mutations arising from stochastic processes (Fig. 1f). The accompanying single-cell resolution ground truth data tracks the complete evolutionary history of each cell, including exact parent-child relationships, cell birth/death events, and precise mutation timing. This comprehensive tracking enables detailed validation of single-cell analysis methods and allows researchers to study cell-to-cell heterogeneity and trace mutation trajectories with high temporal resolution.

In summary, cancerSimCraft provides cancer genome simulation at both clonal and single-cell levels with comprehensive ground truth tracking. These capabilities enable a wide range of applications, from benchmarking bioinformatics tools to investigating complex scientific hypotheses in cancer research.

### Example workflow and validation of clonal-level simulation using cancerSimCraft

To demonstrate cancerSimCraft’s capabilities, we derived an example of a clonal-level simulation workflow with three major stages: simulation design, implementation, and validation (Fig. 2a). For simulation design, we utilized haplotype copy number profiles from a breast cancer sample that was previously analyzed using CHISEL (Zaccaria & Raphael, 2021) (Fig. 2a, left). This dataset, which contains copy number information across 1,448 cells with well-defined clonal structure, provided an ideal basis for evaluating our simulation framework (Fig. 2b). Based on these haplotype copy number profiles and clonal phylogenetic structure, we constructed a simulation blueprint that recapitulates key genomic features in the real data (Fig. 2c). Additionally, we constructed a clonal event tree to visualize the phylogenetic relationships between clones and the genomic alterations that occur along each evolutionary branch (Supplementary Fig. 1b). This blueprint served as the ground truth for subsequent validation of our simulation results. To ensure a comprehensive implementation of cancer genome features, we also incorporated phased SNP information from a high-depth breast cancer dataset and configured clone-specific SNV counts based on literature-derived parameters (Salcedo et al., 2020) (Fig. 1a). Then in the simulation implementation stage, we utilized cancerSimCraft to generate both small-scale (150 cells) and large-scale (3000 cells) datasets (Fig. 2a, middle). Our validation process examined both genome-level features, including chromosome lengths and SNV signals, and read-level features, such as total CNV, haplotype-specific CNV, and SNP/SNV signals, ensuring comprehensive quality assessment of the simulated data (Fig. 2a, right).

**Figure 2.**
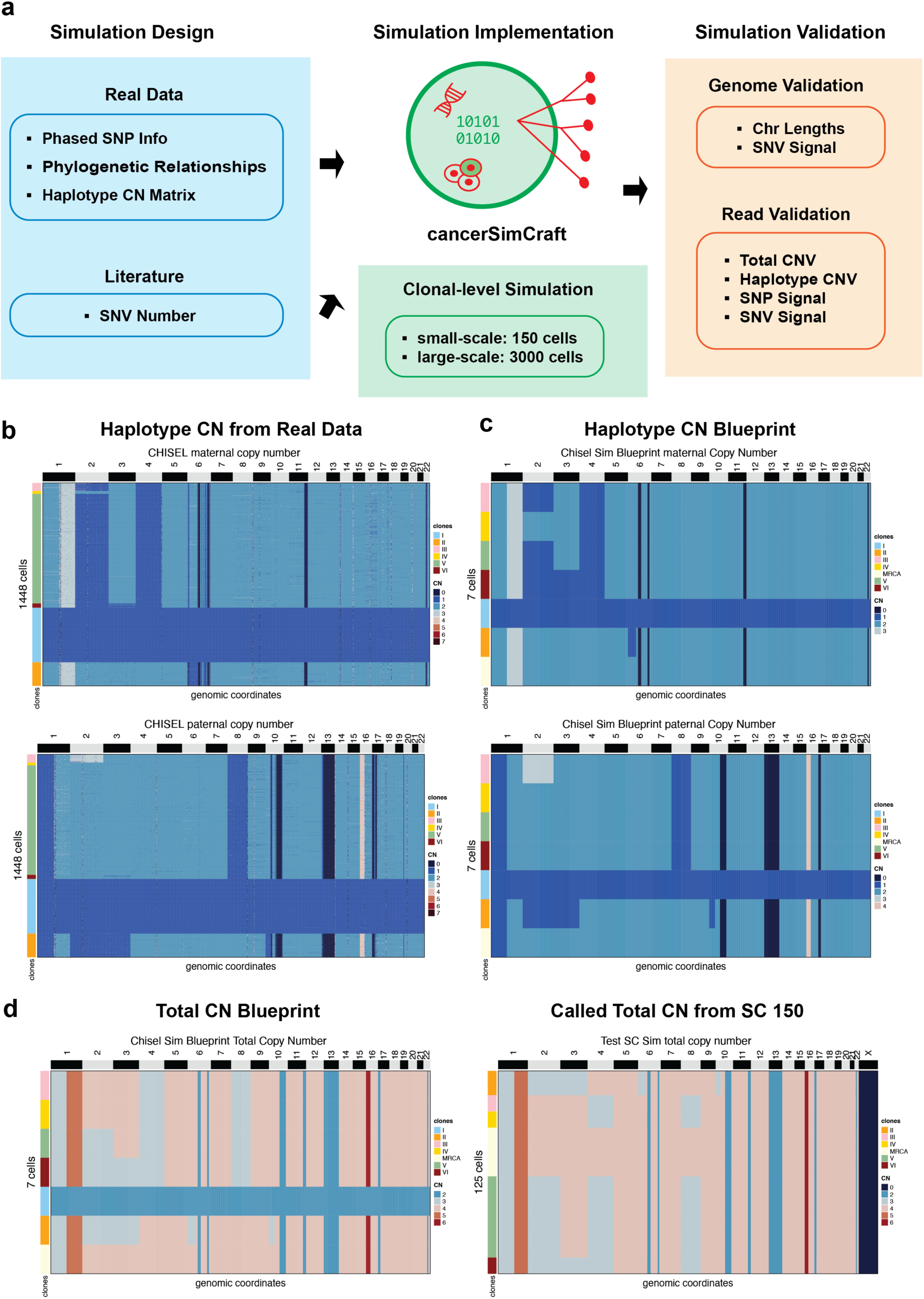
Example workflow and validation of clonal-level simulation using cancerSimCraft. **a,** A schematic of the cancerSimCraft workflow for clonal-level simulation and validation. The workflow is organized into three main stages: (1) Simulation Design – integration of real cancer genomic data and literature information for simulation configuration; (2) Simulation Implementation – running clonal-level simulations using cancerSimCraft, generating both small-scale (150 cells) and large-scale (3000 cells) datasets; and (3) Simulation Validation – assessment of genome-level features (chromosome lengths, SNV signals) and read-level features (total CNV, haplotype CNV, SNP, and SNV signals). **b,** Haplotype-specific copy number matrices from real cancer genome data (Zaccaria & Raphael, 2021) showing maternal (top) and paternal (bottom) copy number states across chromosomes and subclones. **c,** Simulation blueprint derived from real cancer genome data in b. **d,** Validation comparison between blueprint total copy number (left) and total copy number profiles called from simulated data for 150 cells (right, 25 normal cells are eliminated in the heatmap). Comparable results were obtained with 3000 cells (Supplementary Fig. 1a)

Initial validation of our simulation results focused on comparing the copy number profiles between the original blueprint and the simulated data. We observed strong concordance between the total copy number of the blueprint and the 150-cell simulation dataset (Fig. 2d), demonstrating accurate reproduction of complex genomic features such as whole-genome duplications (WGDs) and copy number variations (CNVs). Similar concordance was observed with the larger 3000-cell dataset (Supplementary Fig. 1a), further demonstrating the scalability and robustness of our approach.

To validate cancerSimCraft’s performance at the haplotype level, we examined haplotype-specific alterations and variant patterns in both 150-cell and 3000-cell simulations (Figure 3). Analysis of B-allele frequency (BAF) distributions confirmed the correct incorporation of loss of heterozygosity (LOH) events in chromosome 6 for both 150-cell and 3000-cell datasets (Fig. 3a). In addition, we validated a haplotype-specific duplication event on chromosome 16p. This paternal duplication event was clearly detectable in the large-scale dataset (3000 cells) but was less apparent in the small-scale dataset (150 cells) due to data sparsity (Fig. 3b). This observation highlights the impact of dataset scale on signal detection, underscoring the importance of adequate cell numbers or sequencing depths for accurately detecting certain genomic alterations in single-cell sequencing data.

**Figure 3.**
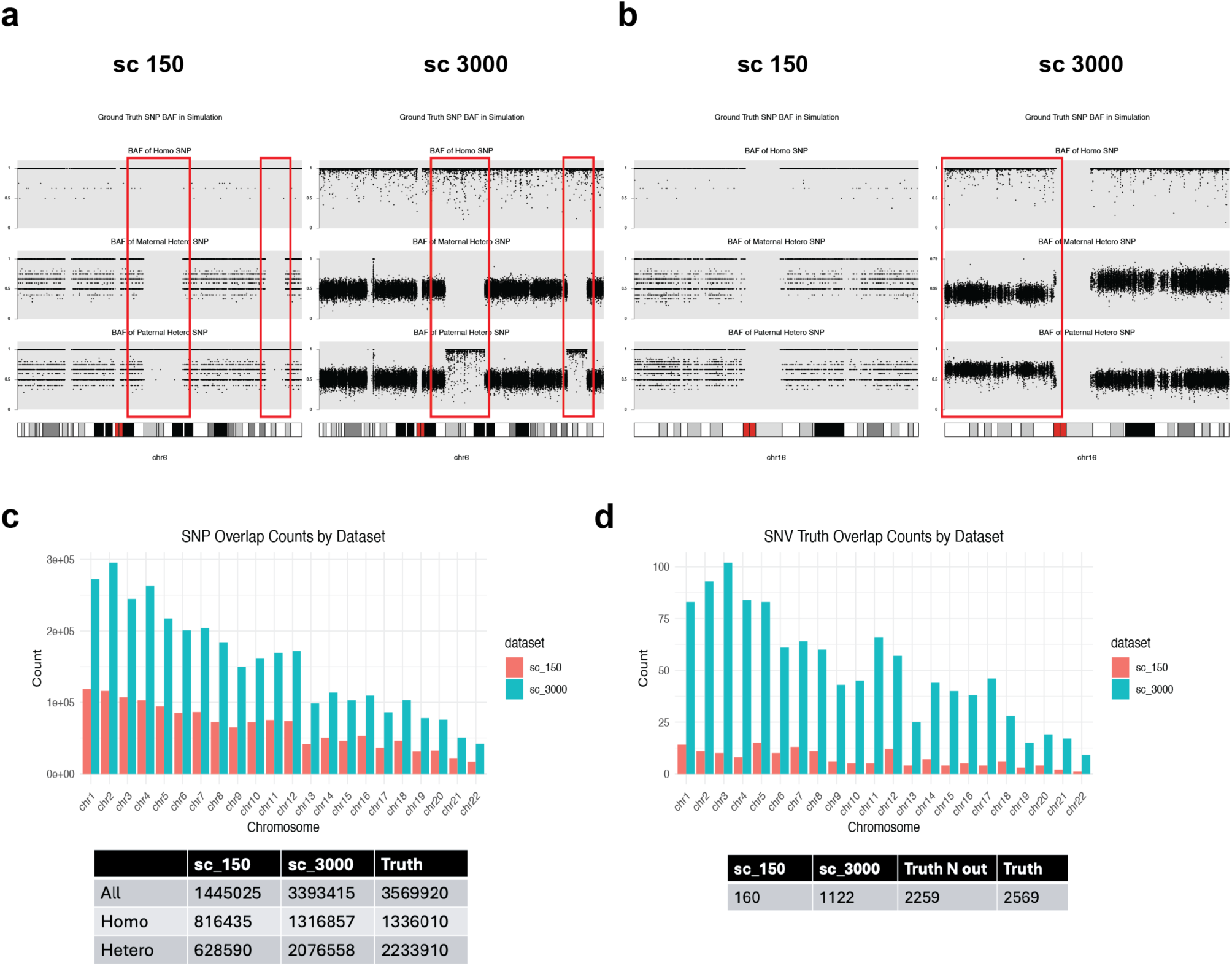
Validation of haplotype-specific events and variant signals in simulated data. **a,** Validation of haplotype-specific loss of heterozygosity (LOH) events on chromosome 6 using SNP allele frequencies. The B-allele frequency (BAF) distributions illustrate LOH at two segments in both small-scale (150 cells) and large-scale (3000 cells) datasets. Red rectangles mark the regions of LOH events. **b,** Validation of a haplotype-specific duplication event on chromosome 16p. The duplication signal at the paternal haplotype becomes clearly visible in the large-scale dataset (3000 cells) but not in the small-scale dataset (150 cells) due to data sparsity. Red rectangles mark the regions of duplication events. **c,** Chromosome-wise comparison of SNP recovery in simulated sequencing data compared to ground truth for both small-scale (150 cells) and large-scale (3000 cells) datasets. Table shows total SNP counts summed over all chromosomes in simulated data (sc_150 and sc_3000) and ground truth, categorized into total SNPs (All), heterozygous SNPs, and homozygous SNPs. **d,** Chromosome-wise comparison of SNV recovery in simulated sequencing data compared to ground truth for both small-scale (150 cells) and large-scale (3000 cells) datasets. Table shows total SNV counts across chromosomes for both simulation scales (sc_150 and sc_3000) compared with all ground truth SNVs (Truth) and theoretically detectable SNVs (Truth_N_out). The “Truth N Out” category represents the theoretically detectable SNVs, excluding regions affected by copy-loss events.

To validate the incorporation of SNPs and SNVs, we first confirmed their complete introduction at the genome level. Then, we examined their recovery in the simulated sequencing data compared to ground truth. Our chromosome-wise comparison analysis revealed successful SNP incorporation in the large-scale simulation, with the 3000-cell dataset recovering 95.1% of ground truth SNPs (3,393,415 vs 3,569,920 total SNPs), including 98.6% of homozygous (1,316,857 vs 1,336,010) and 93.0% of heterozygous SNPs (2,076,558 vs 2,233,910) (Fig. 3c). The smaller dataset (150 cells) showed reduced SNP recovery (1,445,025 total SNPs, 40.5%) due to data sparsity. We then examined SNV recovery across chromosomes, focusing on theoretically detectable mutations that exclude copy-loss regions. Among the 2,259 theoretically detectable SNVs, the 3000-cell and 150-cell datasets recovered 49.7% (1,122 SNVs) and 7.1% (160 SNVs) respectively. The lower recovery rates of SNV reflect both the random sampling nature of sequencing and the inherent sparsity of SNV signals, as SNVs can be either clonal or subclonal, while SNPs are present across all cells in the single-cell dataset (Fig. 3d)

### Systematic benchmarking of computational tools for cancer genome analysis

To evaluate single-cell DNA sequencing analysis methods using cancerSimCraft, we derived a systematic benchmarking framework for evaluating CHISEL’s performance across three key aspects: (1) Technical factors include variations in the dataset size (ranging from 150 to 3000 cells) and the phase list sources. We tested two phase lists: a ground truth phase list (labeled as ‘truth’ in figures) and an inferred phase list from the simulated data using HaplotypeCaller (labeled as ‘haplotypeCaller’ in figures). These diverse technical settings allow us to evaluate CHISEL’s performance under different scenarios of data availability and quality, reflecting challenges encountered in real-world single-cell sequencing applications; (2) Algorithmic factors assess the effect of CHISEL’s three analysis modes: “no-normal,” “pseudo-normal,” and “normal.” These modes represent different approaches to normalization using diploid reference cells. The ‘no-normal’ mode operates without any normal cell reference, while ‘pseudo-normal’ uses diploid cells identified from the scDNA-seq data, and ‘normal’ incorporates high-depth sequenced normal cells. Comparing these modes revealed how CHISEL’s parameter settings influence its accuracy in clone identification and copy number prediction; and (3) Evaluation metrics focus on two key measures: Adjusted Rand Index (ARI) for assessing cancer clone identification and Mean Absolute Error (MAE) for copy number estimation. ARI quantifies the accuracy of CHISEL’s clustering compared to true clonal structure, with higher values indicating better performance. MAE measures the deviation of predicted copy numbers from ground truth for both total and haplotype-specific estimates, with lower values indicating greater accuracy (Fig. 4a).

**Figure 4.**
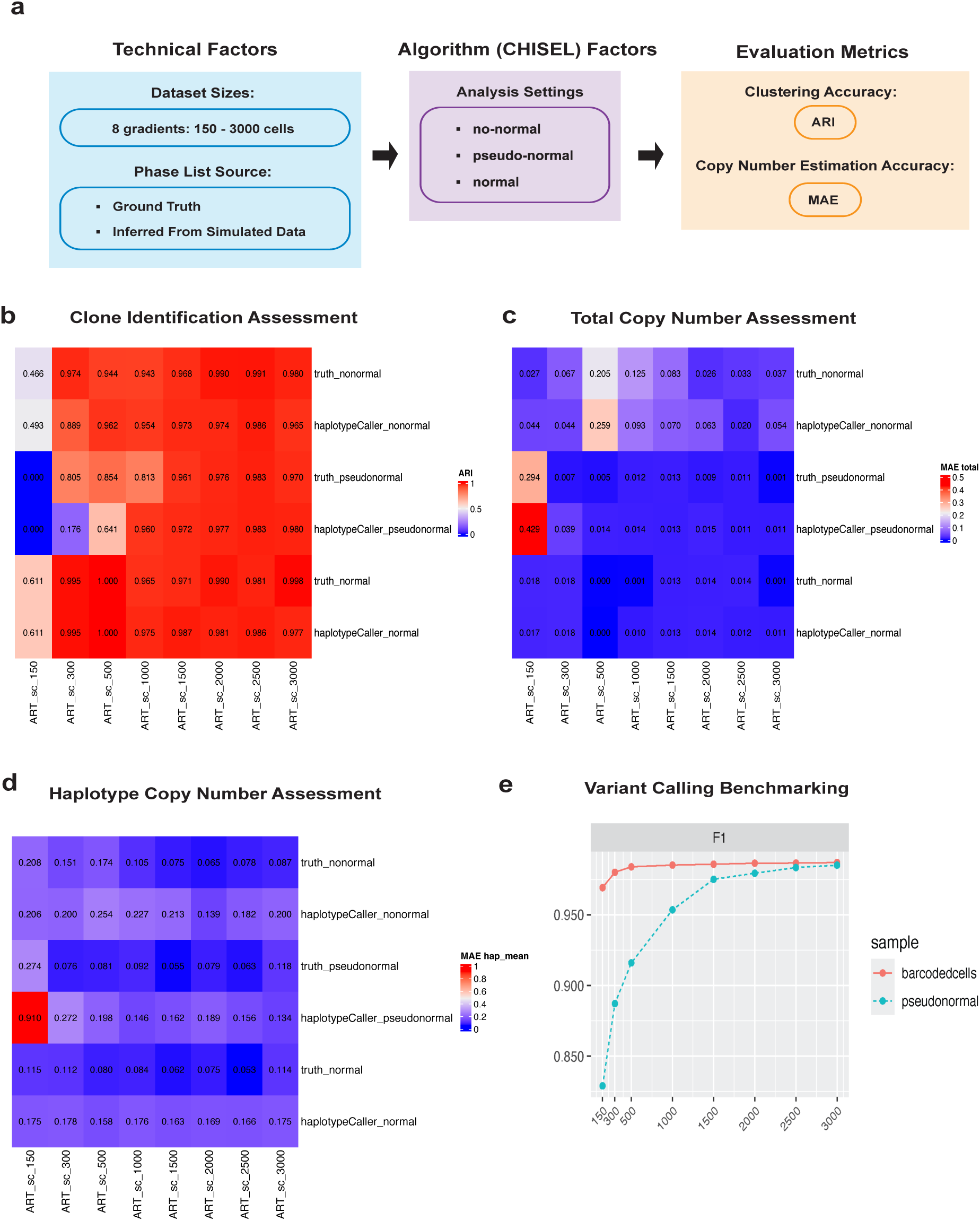
Systematic benchmarking of computational tools for cancer genome analysis. **a,** Framework for evaluating CHISEL’s performance across three key aspects: (1) Technical factors - dataset size (150 to 3000 cells) and phase list source (ground truth vs. inferred); (2) Algorithm factors - CHISEL analysis modes (no-normal, pseudo-normal, and normal); and (3) Evaluation metrics - clone identification accuracy (measured by Adjusted Rand Index, ARI) and copy number estimation accuracy (measured by Mean Absolute Error, MAE) **b,** Assessment of CHISEL’s performance in clone identification using ARI. **c,** Assessment of CHISEL’s performance in total copy number prediction using MAE. **d,** Assessment of CHISEL’s performance in haplotype-specific copy number prediction using MAE. **e,** HaplotypeCaller variant calling performance evaluated using F1 scores under two strategies: using all cells (barcoded cells) versus using detected diploid cells (pseudo-normal).

For clone identification accuracy (Fig. 4b), we observed a clear relationship between dataset size and clustering performance. While small datasets (150-300 cells) showed consistently poor performance across all conditions, datasets with 500-1000 cells demonstrated improved accuracy with only occasional poor performance in specific conditions. Notably, from 1500 cells onwards, clone identification accuracy reached saturation with ARI values exceeding 0.95 across all six benchmarking conditions, regardless of the analysis mode or phase list source. These observations provide practical insights for experimental design, demonstrating that increased dataset size leads to better clone identification performance.

We next assessed total copy number prediction accuracy (Fig. 4c). Similar to clone identification results, we observed performance improvements with increasing dataset size, though saturation occurred earlier at 500 cells, with consistently low MAE values (< 0.015) across most conditions. This early saturation suggests that accurate total copy number estimation may require fewer cells compared to clone identification. However, we observed exceptions in conditions using the ‘nonormal’ mode. Both truth_nonormal and haplotypeCaller_nonormal conditions showed persistently poor performance even with 3000 cells, showing MAE values of 0.037 and 0.054. This consistent underperformance highlights the critical importance of normal cell or pseudo-normal references in accurate copy number estimation, regardless of dataset size.

Haplotype-specific copy number estimation (Fig. 4d and Supplementary Fig. 8a) showed similar dataset size dependence, with performance generally saturating from 1000 cells onwards. Two notable patterns emerged from this analysis. First, across all dataset sizes from 300 to 3000 cells, conditions using truth phase lists consistently achieved lower MAE values compared to those using HaplotypeCaller (McKenna et al., 2010) and Beagle-inferred (Browning et al., 2021) phasing under the same analysis mode. This highlights the impact of phase list accuracy on haplotype copy number inference. Second, we observed that haplotype-specific copy number estimation generally yielded higher MAE values (mean: 0.159) compared to total copy number estimation (mean: 0.048). A detailed comparison between inferred and ground truth copy number profiles in the 3000-cell dataset revealed that while CHISEL accurately identified copy number values, it occasionally assigned copy number states to the incorrect haplotype (Supplementary Fig. 2 and Supplementary Fig. 3). For example, LOH events at chromosomes 6 and 10, which affected different haplotypes in the ground truth (Supplementary Fig. 3), were assigned to the same haplotype in CHISEL’s inference (Supplementary Fig. 2). Similarly, duplications at chromosomes 16 and 1, which affected different haplotypes in the ground truth (Supplementary Fig. 3), were assigned to the same haplotype in the inference (Supplementary Fig. 2). These haplotype assignment discrepancies, rather than errors in copy number values, explain the difference between haplotype-specific and total copy number MAE values. When CHISEL correctly identifies the total copy number (e.g., 3) but swaps the haplotype-specific states (e.g., inferring 2|1 instead of the true 1|2), it produces errors in haplotype-specific assessment while maintaining perfect accuracy in total copy number estimation. This challenge in haplotype assignment stems from the reliance on phased SNP signals (1|0, 0|1), where inaccurate phasing in certain genomic regions can lead to incorrect haplotype assignments. This explains why using ground truth phasing, which avoids such phasing errors, consistently yielded better results.

Finally, we evaluated variant calling performance using HaplotypeCaller (McKenna et al., 2010) under two different strategies: using all cells (barcodedcells) versus using only diploid cells (pseudonormal) (Fig. 4e). The results demonstrated robust variant calling accuracy across both approaches. Even in the more restrictive pseudo-normal strategy, which relies only on reads from diploid cells, F1 scores exceeded 0.9 at 500 cells and improved to above 0.95 at 1000 cells, supported by consistent patterns in precision and recall metrics (Supplementary Fig. 8b). The performance saturation point around 500 - 1000 cells aligns with our observations across this set of benchmarks on clonal assignment (Fig. 4b) and copy number estimation (Fig. 4 c and d).

### Simulation of evolutionary dynamics at the single-cell resolution

The evolution of cancer cell populations involves complex dynamics of clonal expansion, lineage divergence, and mutation accumulation. To capture these dynamics at single-cell resolution, cancerSimCraft employs a continuous-time Markov process to model discrete cellular events (birth, death, transition, and mutation), incorporating the Lotka-Volterra model to account for intercellular competition and population constraints. This approach enables cancerSimCraft to generate cancer genomes that reflect intricate evolutionary processes at single-cell resolution, where each cell’s simulated genome captures both clonal-level alterations and single-cell-specific mutations. The single-cell resolution simulation and ground truth tracking capabilities allow researchers to explore intra-tumor heterogeneity and conduct sophisticated *in silico* experiments in controlled virtual settings.

To demonstrate cancerSimCraft’s ability to model evolutionary dynamics, we simulated three distinct scenarios by modulating birth rates, death rates, and clone carrying capacities: (1) random growth without selective advantage, (2) emergence of a single dominant clone, and (3) emergence of two major clones. Population history plots reveal the distinct dynamics of these scenarios (Fig. 5a). In the random scenario, no single clone dominates the population, leading to a diverse mixture of subclones over time (Fig. 5a, left). In the one major clone scenario, a single clone undergoes significant expansion while other subclones remain in the minority (Fig. 5a, middle). The two major clones scenario shows two distinct clones achieving prominence within the population (Fig. 5a, right).

**Figure 5.**
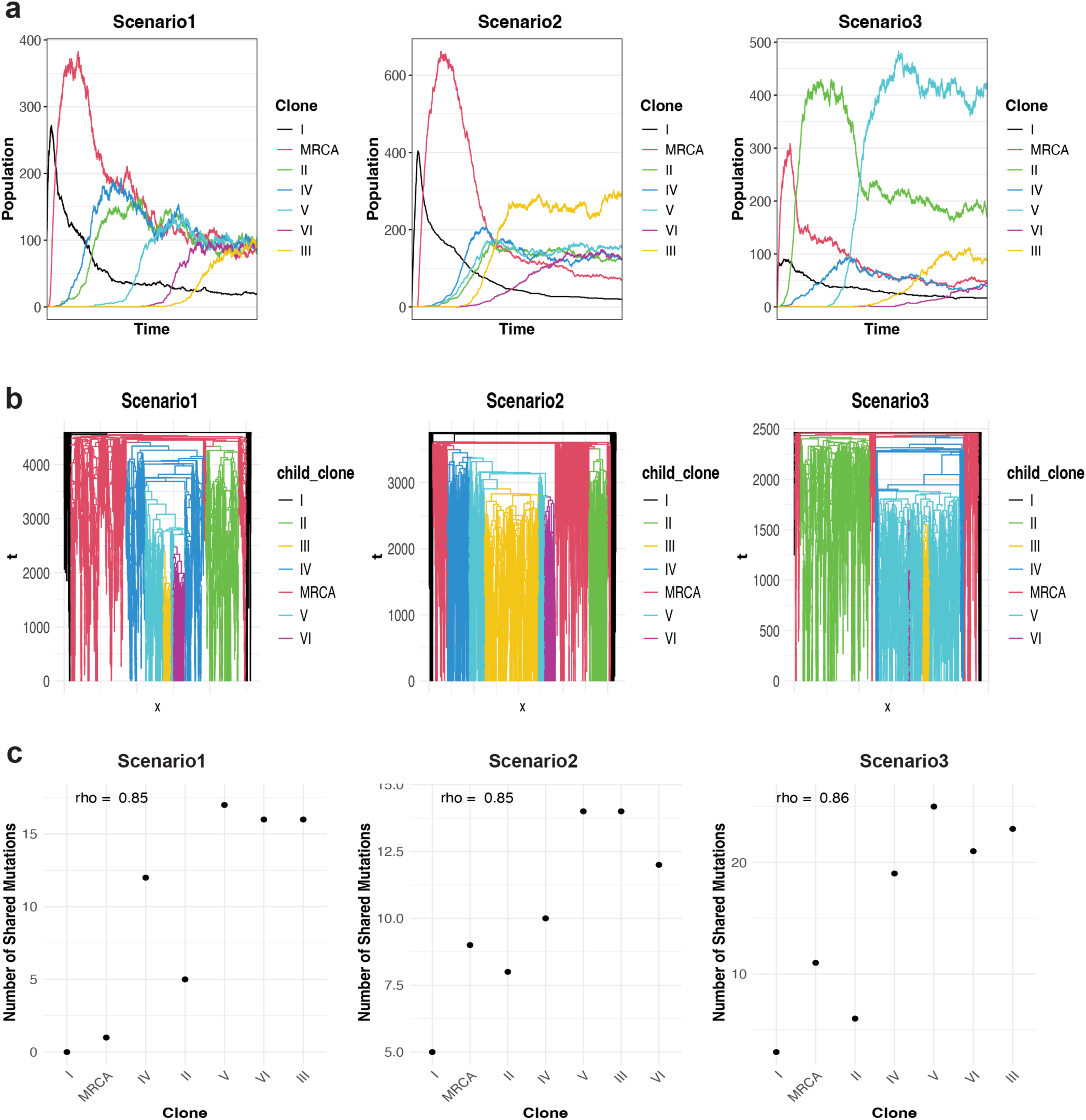
Simulation of evolutionary dynamics at single-cell resolution. **a,** Population dynamics across three evolutionary scenarios: random growth (left), one major clone (middle), and two major clones (right). **b,** Single-cell phylogenetic trees for each scenario, showing lineage relationships and temporal dynamics. **c,** Relationship between clone emergence order (x-axis) and number of shared mutations (y-axis) across scenarios, with Spearman’s correlation (rho) quantifying the association between clonal hierarchy and mutation sharing.

For each scenario, we constructed single-cell phylogenetic trees that reveal lineage relationships and temporal dynamics across all cells (Fig. 5b). Each branch represents an individual cell, with branch lengths indicating the time since divergence from its parent cell. These trees enable precise tracking of evolutionary histories at single-cell resolution, capturing parent-child relationships and the timing of cellular events.

To evaluate reproducibility, we replicated each evolutionary scenario three times. Despite the inherent stochasticity of the Markov process, all replicate simulations exhibited consistent population dynamics when run for sufficient time steps, demonstrating the robustness of our simulation framework (Supplementary Fig. 4 - 6).

We then analyzed the relationship between clone emergence order and mutation sharing across the three evolutionary scenarios. Scatter plots revealed that earlier-emerging clones share fewer mutations while later-emerging clones exhibit larger mutation sharing (Fig. 5c). This pattern was consistent across all replicates, with Spearman’s correlation coefficients exceeding 0.75, demonstrating a robust relationship between clonal hierarchy and mutation-sharing patterns (Supplementary Fig. 7). Such *in silico* experiments are particularly valuable when direct phylogenetic reconstruction is challenging due to data sparsity. While specific mutation signals may be lost in sparse single-cell data, aggregate features such as mutation sharing patterns and burden distributions can still provide meaningful insights.

Together, these results demonstrate the utility of cancerSimCraft in simulating the evolutionary dynamics of cancer at single-cell resolution. By modeling distinct evolutionary scenarios, cancerSimCraft enables us to study how clonal structure, lineage relationships, and mutation burdens evolve over time. These simulations provide a framework for understanding the evolution of cancer subclones, contributing valuable insights into the complex dynamics of tumor heterogeneity and clonal competition.

## Discussion

We present cancerSimCraft, a comprehensive framework that addresses critical needs in cancer genome simulation through several innovative features. First, its multi-resolution simulation capability enables researchers to generate data at both clonal and single-cell levels, providing unprecedented flexibility in studying cancer evolution across different scales. The framework’s ability to capture single-cell dynamics through a continuous Markov process, while simultaneously tracking clonal-level and single-cell-level alterations, makes it particularly valuable for studying intratumor heterogeneity and evolutionary trajectories that are often masked in the sparse and noisy real data. Second, cancerSimCraft’s modular architecture, which separates simulation design from execution, offers practical advantages for research applications. This design allows researchers to efficiently iterate through different simulation parameters without redundant computational processing, facilitating rapid experimental design and hypothesis testing. Third, the framework’s ability to integrate multiple information sources - from real cancer genome data to user-defined hypotheses - enables the creation of highly customized simulations that reflect both empirical patterns and theoretical models. Fourth, cancerSimCraft’s comprehensive ground truth tracking system provides researchers with detailed records of every genomic alteration and cellular event. This complete knowledge of the underlying truth, combined with standard format outputs (fasta and fastq), makes it particularly valuable for developing and validating computational methods in cancer genomics.

Our systematic benchmarking and evolutionary analyses revealed several important insights for method development and experimental design. Through evaluation of CHISEL’s performance, we identified critical dataset size thresholds for different analytical tasks - 500 cells for reliable total copy number estimation and 1500 cells for optimal clone identification. Furthermore, we found that while dataset size is crucial, other factors such as the availability of normal cell references can significantly impact analysis accuracy. Our analysis of haplotype-specific copy number prediction uncovered specific challenges, such as correct copy number values being assigned to incorrect haplotypes due to phasing uncertainties. Together, these findings reveal key technical considerations for both experimental design and analytical strategy in cancer genomics.

Beyond method benchmarking, our evolutionary dynamics simulations demonstrate cancerSimCraft’s utility for theoretical research. Through the continuous-time Markov process, we can generate single-cell datasets that capture cell-to-cell variation and complex population dynamics. Our analysis of different evolutionary scenarios and mutation sharing patterns showed how aggregate mutation features can reveal clonal phylogenetic relationships even in sparse single-cell data. This capability enables researchers to explore hypotheses about tumor evolution that would be difficult to test with real data alone.

Looking ahead, cancerSimCraft’s ability to generate large-scale datasets with comprehensive ground truth labels presents opportunities for machine learning applications in cancer genomics. The simulator’s detailed tracking system provides well-characterized training data for deep learning models, addressing the scarcity of reliable labeled data in the field. These well-characterized datasets generated by cancerSimCraft could advance the development of deep learning models for complex genomic analyses, including mutation timing inference, evolutionary trajectory reconstruction, and tumor heterogeneity characterization.

## Methods

### cancerSimCraft overview

cancerSimCraft provides a systematic framework for simulating cancer genomes at both clonal and single-cell resolution. The framework requires three fundamental inputs: (1) a phylogenetic tree defining evolutionary relationships between cell populations, (2) an event table specifying genomic alterations, and (3) a segment table that divides the genome into predefined regions (bins), where bin sizes can be customized based on analysis requirements. Using these inputs, cancerSimCraft generates cancer genomes through a systematic workflow that combines deterministic rules with stochastic processes.

### Clonal-level simulation

For the clonal-level simulation, cancerSimCraft first constructs a clonal event tree by integrating phylogenetic relationships with subclone-specific genomic alterations. The simulator supports various types of genomic events including single nucleotide variants (SNVs), copy number variations (CNVs), and whole genome duplication (WGD) events. Using a template genome (typically human reference genome) as the starting point, it generates unique genomes for each subclone according to the predefined simulation configuration. Patient-specific variation can be incorporated through an optional SNP table.

### Single-cell population dynamics and simulation

For single-cell simulation, cancerSimCraft implements a continuous-time Markov process that models four discrete cellular events: birth, death, transition, and mutation. The framework uses the Gillespie algorithm to determine event timing and incorporates the Lotka-Volterra model to simulate intercellular competition and population size constraints.

### Output generation and ground truth tracking

cancerSimCraft generates outputs at both clonal and single-cell levels. At the clonal level, it produces standard sequence files (fasta and fastq) along with comprehensive ground truth data covering genomic alterations, clone identities, and phylogenetic relationships. At the single-cell level, each simulated fastq file represents a unique single-cell genome that captures both clonal-level alterations and single-cell-specific mutations arising from stochastic processes. The accompanying single-cell-level ground truth tracks the complete evolutionary history of each cell, including exact parent-child relationships, cell birth/death events, and precise mutation timing.

### Genome update framework

cancerSimCraft implements a systematic approach to update genomic states along the phylogenetic tree. The algorithm traverses the tree in depth-first search (DFS) order, processing genomic events sequentially from parent to child nodes. For each edge in the phylogenetic tree, the simulator processes three types of genomic alterations:

1. Whole Genome Duplication (WGD) events: duplicating all chromosomal segments.
2. Copy Number Variations (CNVs): updating copy numbers at chromosome-arm level.
3. Focal alterations (sub_CNVs): handling copy number changes at specific genomic regions.

For each node, the simulator keeps track of copy number states and chromosome lengths, which are updated according to the specific event types encountered along each edge of the phylogenetic tree. This hierarchical updating process ensures accurate propagation of genomic alterations through the predefined evolutionary trajectory while maintaining the integrity of haplotype-specific information.

### SNP introduction

Before genome and read simulation, cancerSimCraft provides an option to introduce SNPs into the template genome through a systematic process. The process requires phased VCF (Variant Call Format) data as input and only considers single-nucleotide-length variants. SNP insertion handles maternal and paternal haplotypes separately, introducing homozygous (1|1) SNPs into both haplotypes and heterozygous SNPs (0|1 or 1|0) into their respective haplotypes. Following insertion, a quality control step verifies that the alternate alleles in the original VCF match the extracted nucleotides in the modified genome. The modified genome sequences serve as templates for subsequent read simulation and analysis steps.

### Genome and read data synthesis

Our simulation framework implements a hierarchical approach that generates both genome sequences and sequencing reads. At the clonal level, we traverse the phylogenetic tree in depth-first search (DFS) order and process each clone’s genome at the haplotype level. Copy number alterations and clone-specific mutations are applied separately to maternal and paternal haplotypes. For each clone, we construct its founder genome by incorporating all inherited alterations from ancestor clones along with its unique genomic changes.

For single-cell genome synthesis, we introduce cell-specific mutations into the founder genomes. Mutations are applied separately to maternal and paternal haplotypes, with each mutation assigned a unique identifier to track its timing and inheritance. We then validate the accuracy of mutation placement through systematic quality checks.

For read simulation, we employ the ART (MountRainier version) (Huang et al., 2012) to generate realistic single-cell sequencing data. For each cell, we combine its maternal and paternal haplotype genomes into a single FASTA file. In the simulation configuration, users can specify key sequencing parameters including read length (default 150bp), sequencing depth (default 30x), and sequencing mode (single-end or paired-end). To optimize performance, the read simulation is parallelized across multiple cores.

### Simulation Design

To establish simulation configurations that reflect real cancer genomes, we extracted copy number profiles from single-cell DNA sequencing data analyzed by CHISEL (Zaccaria & Raphael, 2021). We first identified genomic segments with consistent copy number patterns using a sliding window approach. For each chromosome and haplotype, we calculated Spearman correlations between copy number profiles across genomic windows. Adjacent regions were clustered using a dynamic window extension algorithm: starting with a minimum cluster size, windows were iteratively extended if the average pairwise correlation within the window remained above a threshold (0.85). This approach enabled identification of contiguous genomic segments affected by the same copy number events.

To provide biological context, we mapped these computationally identified regions to cytogenetic bands based on UCSC genome browser annotations (hg19/hg38). Chromosome arms, defined by centromere positions from the UCSC annotation, serve as the primary genomic segments in our framework. Within these chromosome arms, we derived smaller segments to capture focal CNV events. Each segment was assigned a unique identifier to track its evolution throughout the simulation. Using this segmentation approach, we captured key genomic events observed in the real data, including whole genome duplications, loss of heterozygosity (LOH), and copy number alterations at both chromosome-arm and focal levels. These genomic patterns formed the basis for our simulation configurations.

### Mathematical formulation of evolutionary dynamics

#### Initial population

We denote N_i_(t) as the population of cells belonging to clone *i* at time *t*. The initial population size is given by {*N*_1_(0), *N*_2_(0), …, *N_K_*(0)}, where *t* = 0 and *K* is the total number of clones in the dataset.

#### Birth and death rates

Let *N_i_*(*t*) be the population size of clone *i* at time *t*, the intrinsic birth rate of clone *i* is *b_i_* and the intrinsic death rate is *d_i_*. The birth rate is adjusted by carrying capacity *C_i_*:

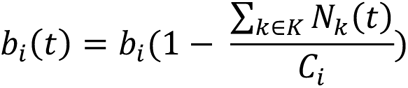

Death rate is proportional to the current population size:

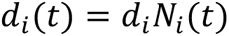

#### Transition rates

Transitions between clones follow specific parent-child relationships defined in set *T*. Each allowed transition (*p*, *c*) ∈ *T* has a base rate *T_pc_*, and the parent clone’s population *N_p_*(*t*) affects all its possible transitions. The total transition rate at time *t* sums over all allowed transitions:

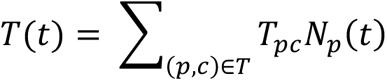

#### The total rate

The total rate *R*(*t*) of all possible events (birth, death, transition) is the sum of individual rates:

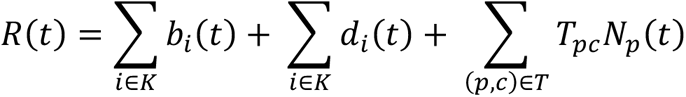

Event timing is determined using the Gillespie algorithm, where the time until the next event follows an exponential distribution with rate parameter *R*(*t*).

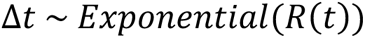

The next event type (birth, death, transition) is chosen based on their relative rates.

#### Mutation process

During each birth event, mutations occur with a specified probability (mutation rate) and are assigned to one of the daughter cells. The mutation simulation follows a two-stage process. First, the mutation location is determined by:

1. Random selection of haplotype
2. Chromosome selection weighted by chromosome lengths
3. Random position selection within the chosen chromosome

Second, for nucleotide changes, we identify the original nucleotide from the reference genome and sample an alternative nucleotide based on a transition probability matrix. For positions in copy-loss regions, both original and alternative nucleotides are set to ‘N’. For each mutation, we record its full characteristics: clone identity, cell index, affected haplotype, chromosome, position, original and alternative nucleotides, and timing.

### Evaluation metrics

We evaluated method performance using two standard metrics. Mean Absolute Error (MAE) measures copy number prediction accuracy by calculating the average absolute difference between predicted and true copy numbers, with lower values indicating better accuracy. Adjusted Rand Index (ARI) assesses clone identification accuracy by quantifying the agreement between predicted and true cell clustering assignments. ARI values range from -1 to 1, where 1 indicates perfect agreement and 0 represents random assignment.

### Processing of simulated sequencing data

Raw sequencing data processing followed a sequential pipeline: (1) alignment to hg19 reference genome using Bowtie2 (v2.3.5.1) (Langmead & Salzberg, 2012), (2) conversion to BAM format and coordinate sorting using SAMtools (v1.9) (Danecek et al., 2021), (3) BAM indexing, and (4) duplicate marking using Sambamba (v0.8.0) (Tarasov et al., n.d.). The pipeline was implemented using Snakemake (v7.8.5) (Köster & Rahmann, 2012) workflow management system to ensure reproducibility and parallel processing efficiency.

### Variant calling and phasing

Variant analysis followed GATK4 best practices (McKenna et al., 2010). Variants were called per-chromosome using HaplotypeCaller, followed by SNP selection and quality filtration. The pipeline employed bcftools (v1.16) (Danecek et al., 2021) for VCF manipulation and merging, with final outputs indexed using tabix (v0.2.6) (Li, 2011). Haplotype phasing was performed using Beagle (v5.4) (Browning et al., 2021) with the 1000 Genomes Phase 3 reference panel (Auton et al., 2015). All steps were orchestrated using Snakemake (v7.32.4) (Köster & Rahmann, 2012) workflow management system to ensure reproducibility.

### Copy number analysis

Total copy number states were determined using Copykit (v0.1.0) (Minussi et al., 2022). Haplotype-specific copy number analysis was performed using CHISEL (v1.1.3) (Zaccaria & Raphael, 2021) under three configurations: (1) no-normal mode without reference cells, (2) pseudo-normal mode using inferred diploid cells from scDNA-seq data, and (3) normal mode with high-depth sequenced normal reference. Each mode utilized corresponding phased SNP information, with analyses performed using multi-threading and fixed random seed for reproducibility.

## Code availability

cancerSimCraft is implemented in R and is freely available as an open-source software under MIT license at https://github.com/haijingjin/cancerSimCraft.

## Author contributions

H.J. conceived and developed the cancerSimCraft, designed and implemented the algorithms, performed all analyses, and wrote the manuscript. N.N. and K.C. supervised the project and provided feedback on the manuscript.

## Acknowledgement

H.J. is a TRIUMPH Fellow in the CPRIT Training Program (RP210028). We thank the TRIUMPH program director, Dr. Khandan Keyomarsi, and program staff for their administrative support and mentorship. We are grateful to H.J.’s TRIUMPH committee members - Drs. Peter Van Loo, Linghua Wang, Anna Aparicio, and Bora Lim - for their thoughtful guidance and support.

---

**Supplementary Fig. 1.**
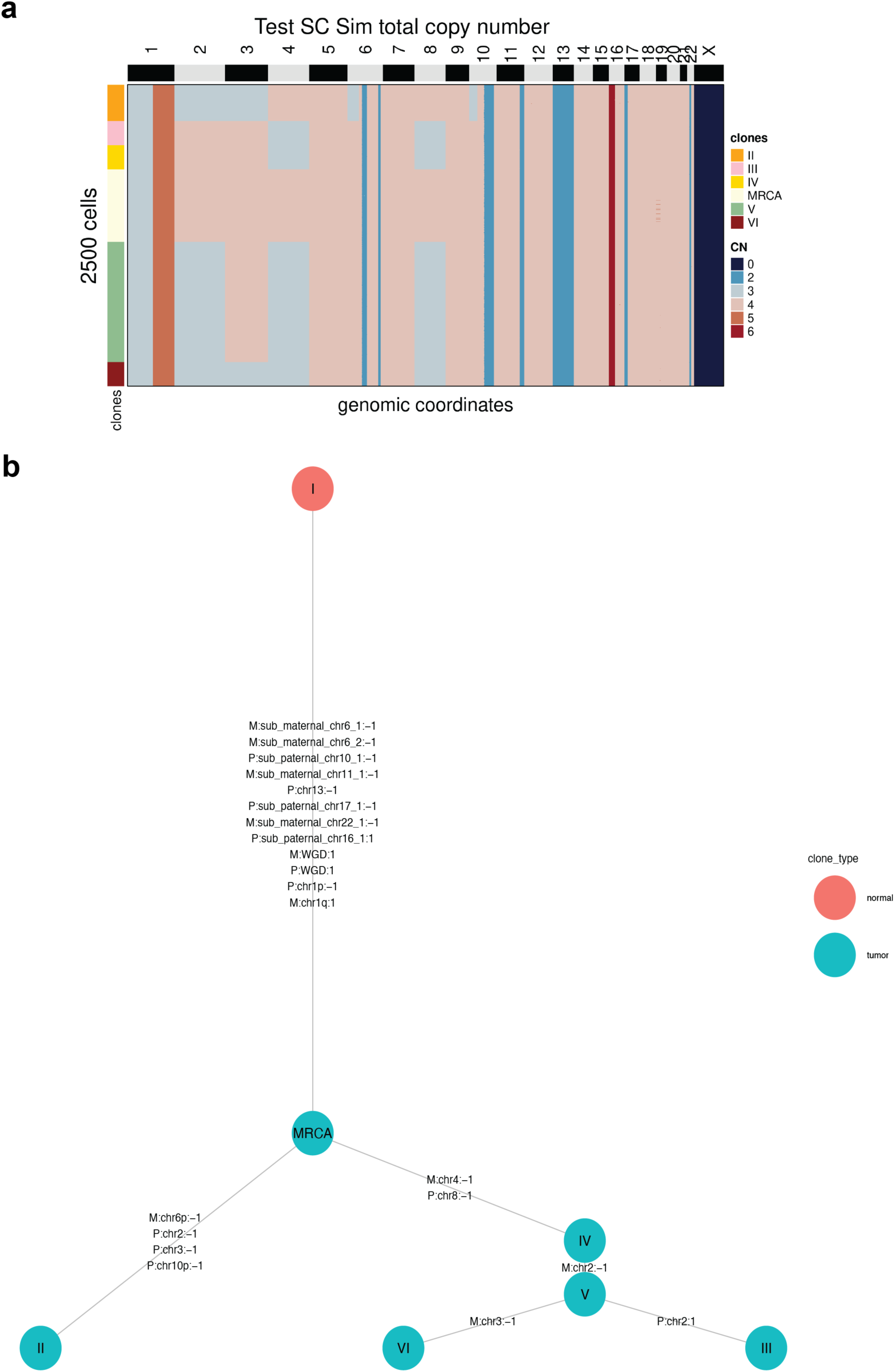
**a,** Total copy number profile called from 3000-cell simulated data, demonstrating robust reproduction of blueprint patterns at larger dataset size. **b,** Clonal event tree showing evolutionary relationships between clones, with edges annotated in the format of haplotype:affected_segment:copy number change. M and P denote maternal and paternal haplotypes. Clone I (red) represents the normal diploid cell state.

**Supplementary Fig. 2.**
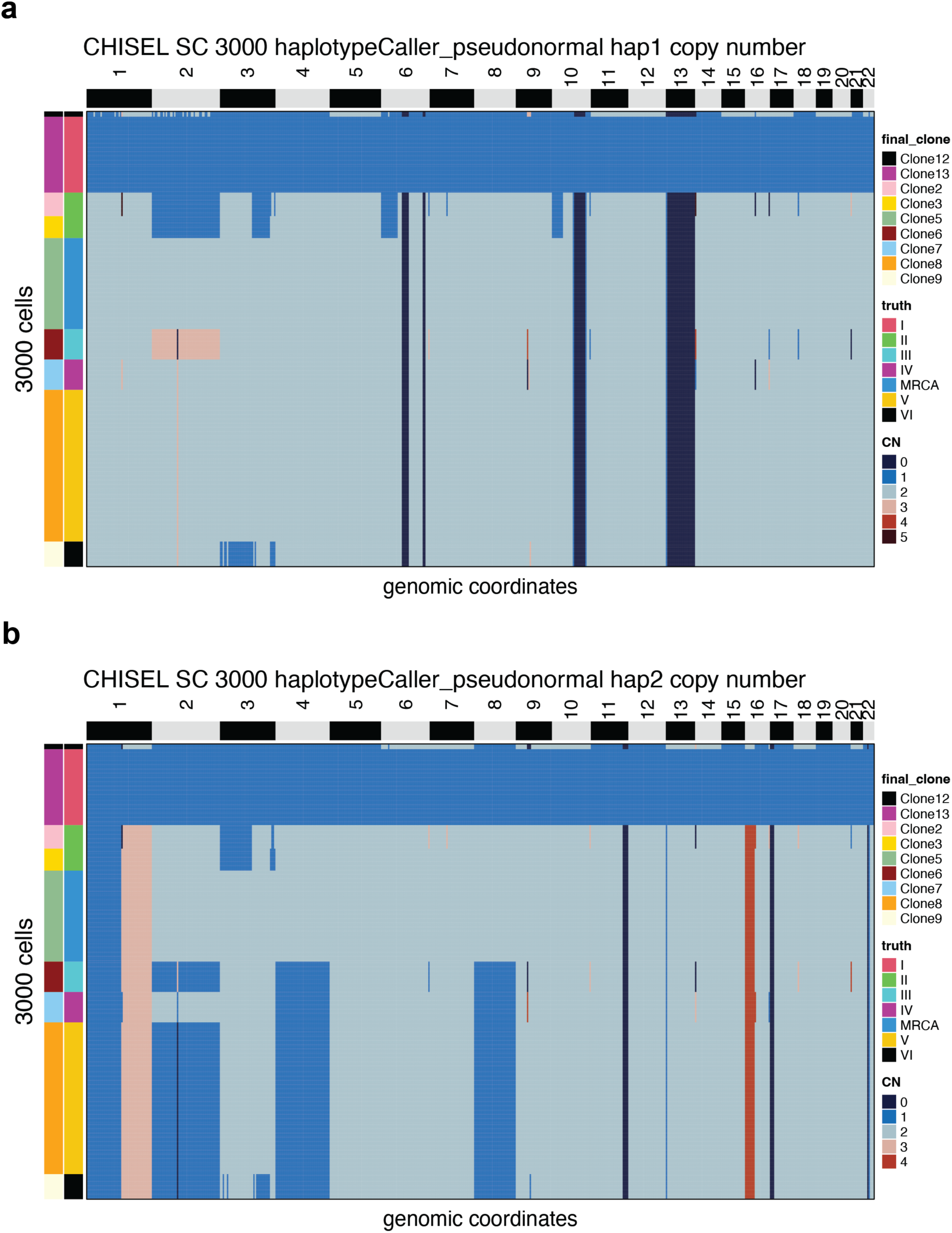
Haplotype-specific copy number profiles inferred by CHISEL from 3000-cell simulation data (haplotypeCaller_pseudonormal setting).

**Supplementary Fig. 3.**
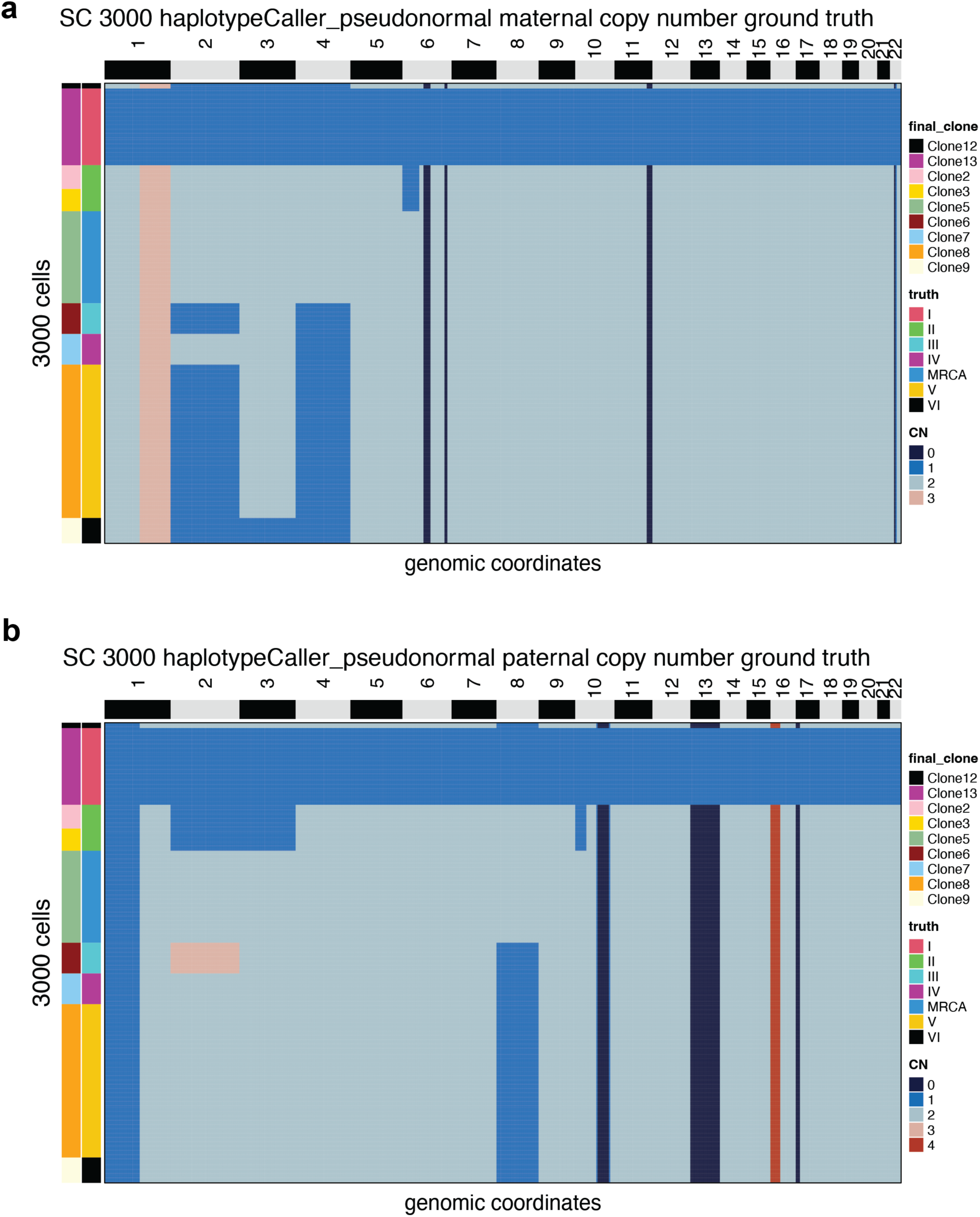
ground truth haplotype-specific copy number profiles for the 3000-cell simulation data, corresponding to CHISEL analysis in Supplementary Fig. 2.

**Supplementary Fig. 4.**
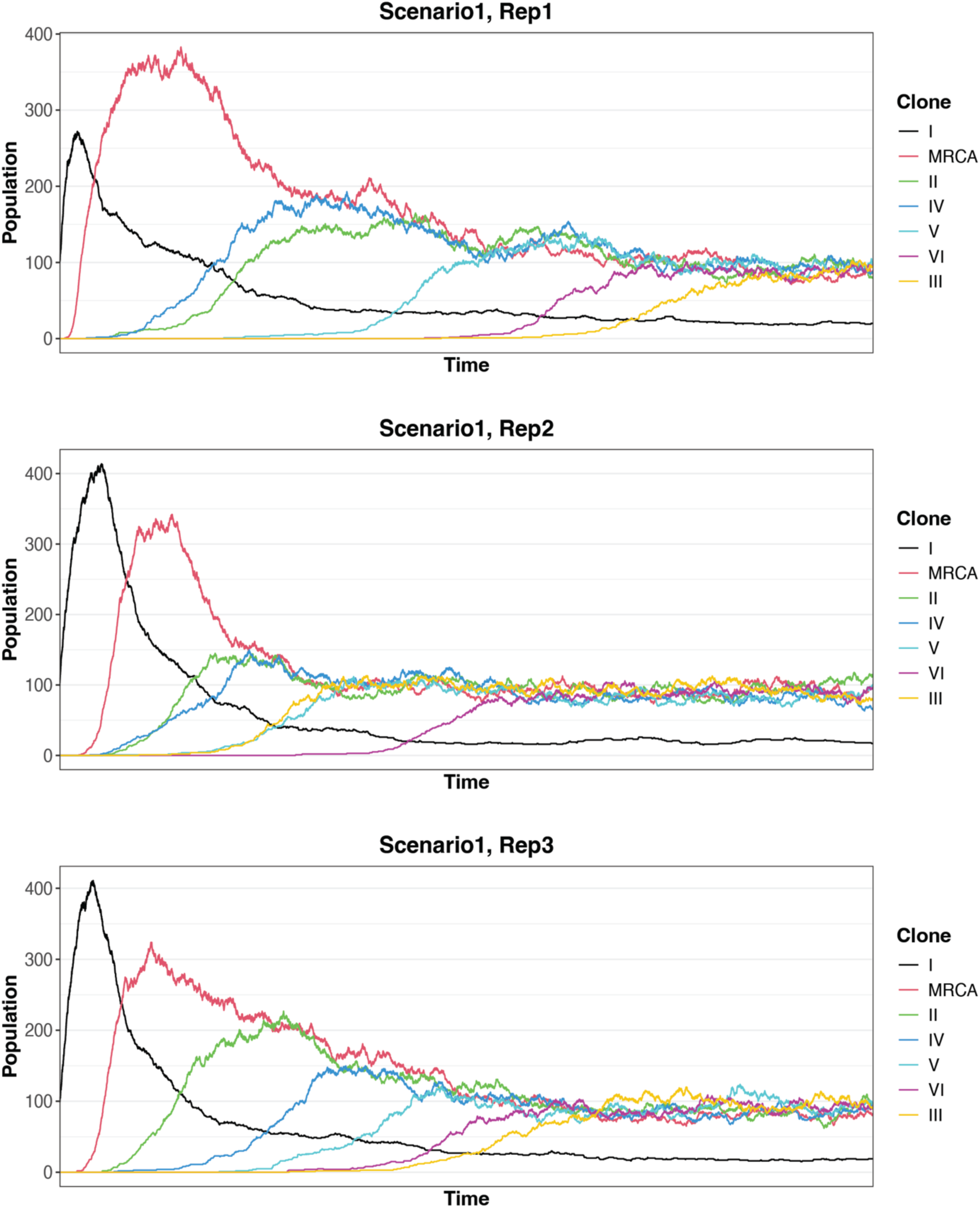
Population dynamics across three replicate simulations of the random growth scenario (scenario 1).

**Supplementary Fig. 5.**
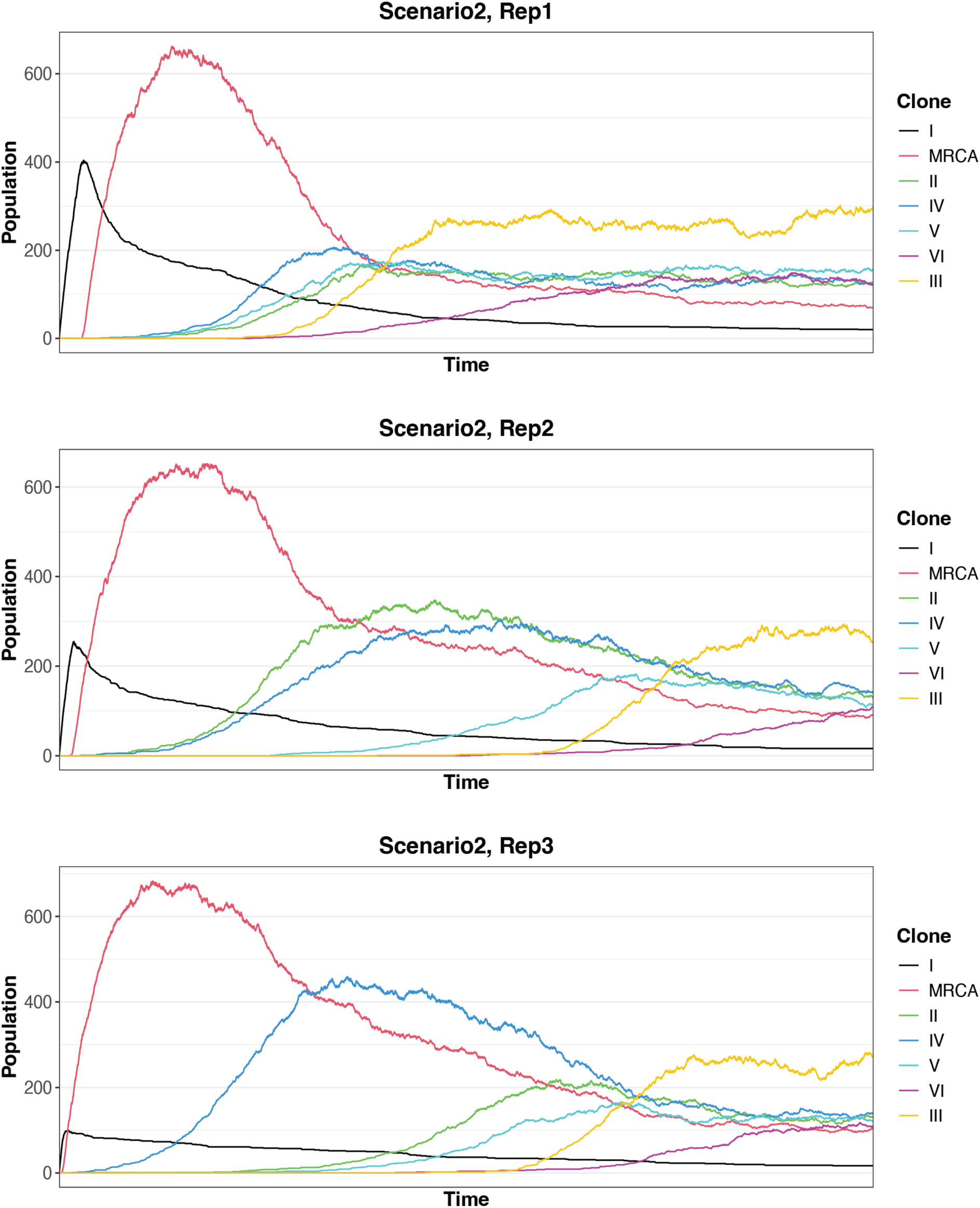
Population dynamics across three replicate simulations of the one major clone scenario (scenario 2).

**Supplementary Fig. 6.**
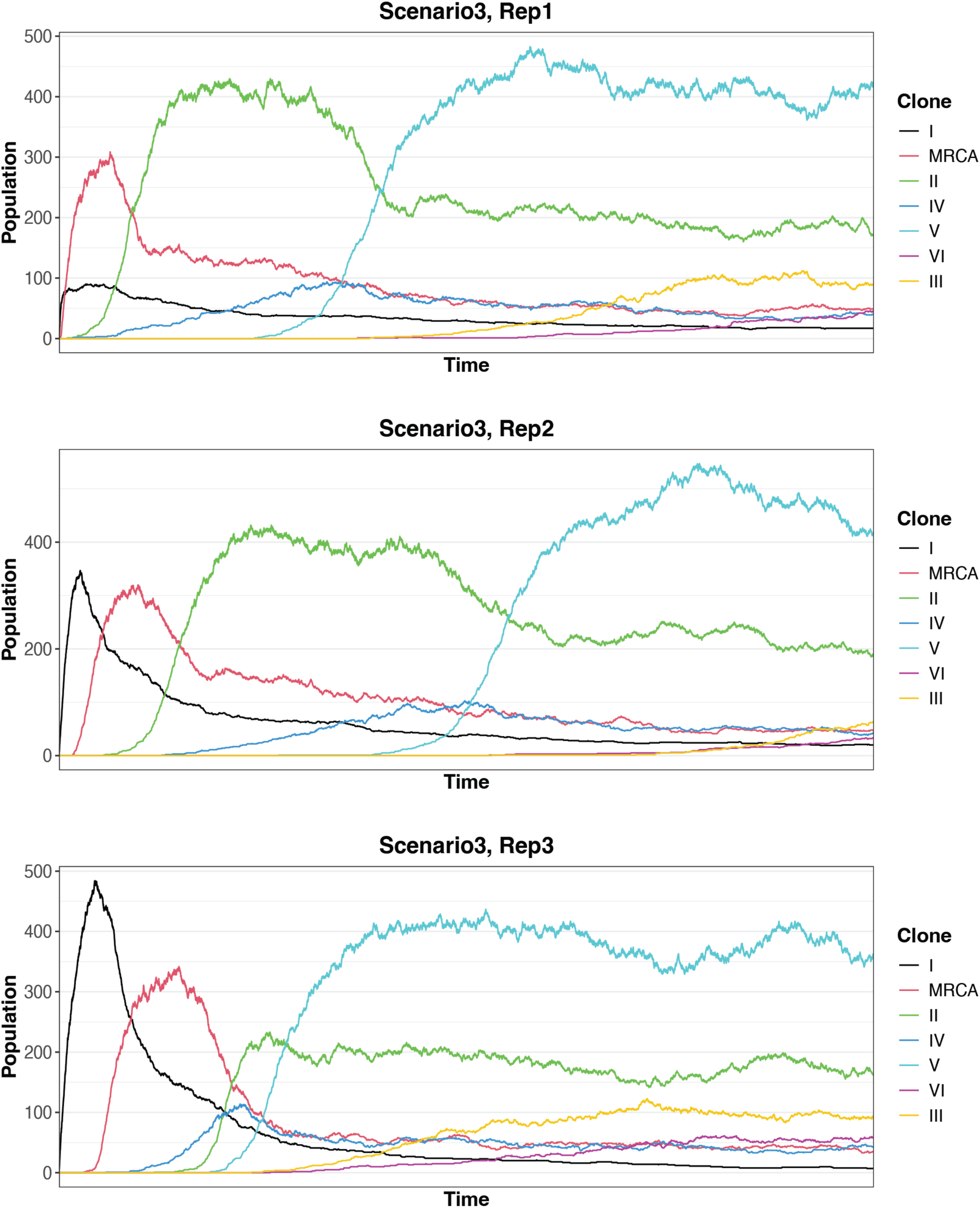
Population dynamics across three replicate simulations of the two major clones scenario (scenario 3).

**Supplementary Fig. 7.**
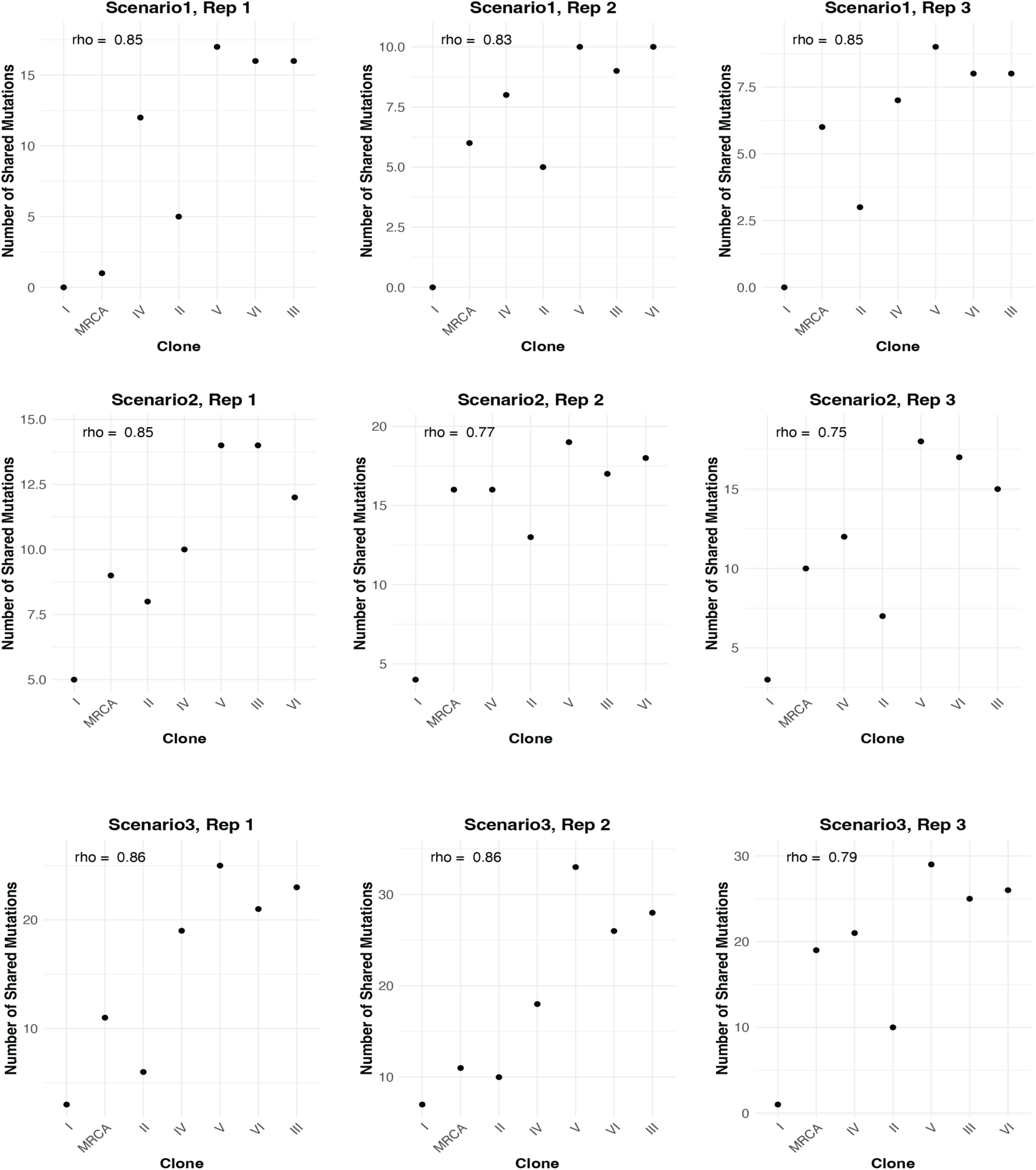
Scatter plots showing the correlation between clone emergence order (x-axis) and number of shared mutations (y-axis) for three replicates of each evolutionary scenario: random growth (scenario 1, top), one major clone (scenario 2, middle), and two major clones (scenario 3, bottom). Spearman’s correlation coefficients (rho) demonstrate consistent relationships across replicates.

**Supplementary Fig. 8.**
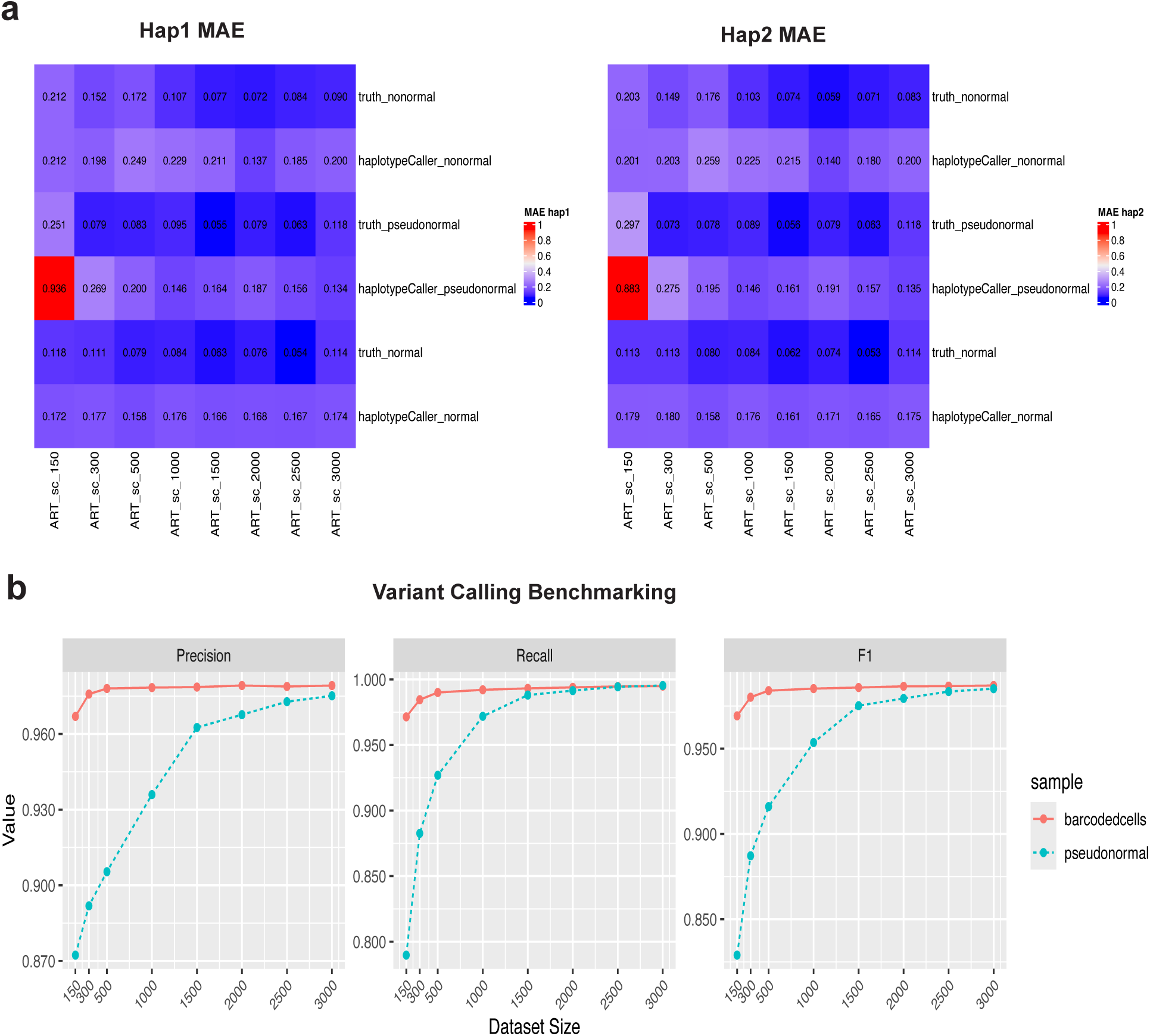
**a,** CHISEL’s haplotype-specific copy number prediction accuracy shown separately for haplotype 1 (left) and haplotype 2 (right), measured by MAE. The mean of these values is shown in Fig. 4d. **b,** Detailed evaluation of HaplotypeCaller variant calling performance showing precision (left), recall (middle), and F1 scores (right) under two strategies: using all cells (barcoded cells) versus using detected diploid cells (pseudo-normal). F1 scores are also shown in Fig. 4e.

## Notes

### Competing Interest Statement

The authors have declared no competing interest.

